# Transcriptional differences between the two host strains of *Spodoptera frugiperda* (Lepidoptera: Noctuidae)

**DOI:** 10.1101/263186

**Authors:** Marion Orsucci, Yves Moné, Philippe Audiot, Sylvie Gimenez, Sandra Nhim, Rima Naït-Saïdi, Marie Frayssinet, Guillaume Dumont, Jean-Paul Boudon, Marin Vabre, Stéphanie Rialle, Rachid Koual, Gael J. Kergoat, Rodney N. Nagoshi, Robert L. Meagher, Emmanuelle d’Alençon, Nicolas Nègre

**Affiliations:** DGIMI, Univ Montpellier, INRAE, Montpellier, France; CBGP, INRAE, IRD, CIRAD, Institut Agro, Univ Montpellier, Montpellier, France; UE Diascope, INRAE, 34130 Mauguio, France; MGX, Univ Montpellier, CNRS, INSERM, Montpellier, France; CMAVE USDA-ARS, Gainesville, Florida, USA; Department of Plant Biology, Swedish University of Agricultural Sciences, Uppsala BioCenter and Linnean Centre for Plant Biology, 75007 Uppsala, Sweden; Department of Microbiology & Immunology, Center for Advanced Microbial Processing, Drexel University College of Medicine, 245 N. 15th Street, Philadelphia, PA, 19102, USA

**Keywords:** Life histoy traits, transcriptomic, mitochondrial variation, species divergence, fall armyworm

## Abstract

*Spodoptera frugiperda*, the fall armyworm (FAW), is an important agricultural pest in the Americas and an emerging pest in sub-Saharan Africa, India, East-Asia and Australia, causing damage to major crops such as corn, sorghum and soybean. While FAW larvae are considered polyphagous, differences in diet preference have been described between two genetic variants: the corn strain (sf-C) and the rice strain (sf-R). These two strains are sometimes considered as distinct species, raising the hypothesis that ost plant specialization might have driven their divergence. To test this hypothesis, we irst performed controlled reciprocal transplant (RT) experiments to address the impact of plant diet on several traits linked to the fitness of the sf-C and sf-R strains. The phenotypical data suggest that sf-C is specialized to corn. We then used RNA-Se to identify constitutive transcriptional differences between strains, regardless of diet, in laboratory as well as in natural populations. We found that variations in mitochon rial transcription levels are among the most substantial and consistent differences between the two strains. Since mitochondrial genotypes also vary between the strains, we believe the mitochondria may have a significant role in driving strain divergence.

## Introduction

The relatively recent development of agroecosystems modified ecological niches in many ways (O’Brien and Laland 2012). First and foremost, artificial selection used by early farmers in south-west Asia as of 10,000 years ago to improve their crops, elicited the rapid apparition of new domesticated varieties in the biosphere (Zohary, Hopf, and Weiss 2012). Whilst being selected for human favored traits, cultivated plants concomitantly lost or gained additional properties and thus plant-interacting organisms were prone to exploit these new niches. For example, some phytophagous insects were able to adapt to cultivated plants and, with the intensification of production based on monoculture activities, these insects eventually became agricultural pests. This adaptation to agricultural plants provides an interesting model system to observe evolution at a relatively small time-scale and assess the genetic changes that may promote speciation in relation to environmental changes (Yoder et al. 2010).

*Spodoptera frugiperda* (J.E. Smith) (Lepidoptera: Noctuidae: Noctuinae), also known as the fall armyworm (FAW), constitutes a good model to study adaptation of phytophagous insects to agricultural plants. Its native distribution range spans a vast amount of the Americas from Brazil to Canada (Pogue 2002). The FAW has no winter diapause (Sparks 1979) and its wintering range is constrained to warmer regions such as southern Florida and southern Texas in the United States (Nagoshi and Meagher 2004). In 2016 it became invasive on the African continent where massive crop damages have been observed across sub-Saharan Africa in less than a year (Goergen et al. 2016; Jeger et al. 2017). It has since been reported in India, South-East Asia, China in 2019 and Australia in 2020 (see the most recent report maps at https://www.cabi.org/isc/fallarmyworm), surely becoming a world-wide menace.

The FAW is a polyphagous species, being documented on over 353 plants from 76 different plant families (Montezano et al. 2018). However, using allozymes electrophoresis monitoring, a significant genetic heterogeneity has been observed in FAW populations that was associated with feeding preferences (Pashley et al. 1985; Pashley 1986). One genetic haplotype was mostly found on corn (*Zea mays*), sorghum (*Sorghum* spp.) and cotton (*Gossypium* spp.) and was named the corn strain (sf-C). Another haplotype was found associated to individuals collected on smaller grasses such as turf, Bermuda (*Cynodon dactylon*) grasses and rice (*Oryza* spp.), and has been named the rice strain (sf-R) (Pashley 1988). Subsequent studies have confirmed these genetic differences on markers such as the mitochondrial gene cytochrome oxidase c subunit I (COI) (Lu and Adang 1996; Meagher and Gallo-Meagher 2003; Nagoshi et al. 2006; Machado et al. 2008), but also nuclear loci, such as the sex-linked FR1 repeat element (Nagoshi and Meagher 2003a; Nagoshi and Meagher 2003b; Lu et al. 1994) and the Z chromosome-linked *Tpi* gene (Nagoshi 2010). Phylogenetic analyses based on COI only (Dumas et al. 2015a; Le Ru et al. 2018) or on several mitochondrial and nuclear markers (Kergoat et al. 2012; Le Ru et al. 2018) showed that sf-C and sf-R separate in two distinct clades that could represent incipient species. While some degree of hybridization has been reported in field samples (Prowell, McMichael, and Silvain 2004; Nagoshi and Meagher 2003a; Nagoshi et al. 2006; Machado et al. 2008), it has also been shown that pre- and post-zygotic reproductive isolation mechanisms exist between the strains (Groot et al. 2010), with a loss of viability of the hybrids (Dumas et al. 2015b; Kost et al. 2016). Differences in reproductive behavior were also documented, such as the timing of mating being shifted earlier in the night for sf-C compared to sf-R (Schöfl, Heckel, and Groot 2009; Groot et al. 2010; Pashley and Martin 1987; Pashley, Hammond, and Hardy 1992). In order to detect post-zygotic reproductive barriers, many studies tried to quantify the impact of the diet on the general fitness of the FAW larvae (Groot et al. 2010; Roy et al. 2016; Meagher et al. 2004; Silva-Brandão et al. 2017; Pashley 1988; Whitford et al. 1988). The results of these studies are sometimes contrasted but seem to agree about a better performance of sf-C on corn indicating that sf-C might be specializing to corn (Groot et al. 2010).

In order to understand if plant adaptation is indeed at the origin of the differences between the strains, we first conducted phenotypical experiments in the context of oviposition choice (OV) to different plants and of a reciprocal transplant (RT) during which we surveyed fitness associated traits (also called Life History Traits or LHT; Stearns 2012) to estimate the preference-performance of both strains. In parallel, we performed RNA-Seq experiments to search for genes constitutively differently transcribed between strains, in laboratory as well as in natural populations, that could indicate which selective pressure led to strains divergence. Surprisingly, we identified a major difference in the transcription of the mitochondrial genome. Since mitochondrial genotypes are also the main genetic variation between the strains, we propose that the mitochondrial genome was the primary target of selection between the two strains.

## Results and discussion

### Difference in oviposition choice between sf-C and sf-R

Under the preference-performance hypothesis, the choice of host plants by adult females to lay their eggs should reflect the host plants on which the larval performance is higher (Thompson 1988; Jaenike 1990; Gripenberg et al. 2010; Clark, Hartley, and Johnson 2011). We conducted an oviposition choice experiment where *S. frugiperda* adult females of each strain (sf-C or sf-R) were set free to lay eggs in a cage containing either their preferred host plant, their alternative host plant (“no-choice” trial) or both (“choice” trial). We recorded the number of egg masses laid by females in each cage, depending on the substrate (the plant type or the cage net). Analysis by a generalized linear model (see **Methods**) showed that the interaction between the strain and the experimental factors was not significant (LRT, F = 1.29, df = 2, *P* = 0.1644). Indeed, we found that the number of egg masses laid by females (Mean fertility) was similar between trials (LRT, F = 0.29, df = 2, *P* = 0.75) but significantly different according to the strain (LRT, F = 24.73, df = 1, *P* < 0.001). Effectively, sf-C laid almost double the number of egg masses than sf-R (Mean fertility of 3.89 for sf-C against 2.06 for sf-R across all trials; **Fig. S1A**). When we analyzed the percentage of egg masses hatching within each trial, we observed no significant difference between strains (LRT, χ^2^ = 0.17, df = 1, *P* = 0.68) or laying sites (LRT, χ^2^ = 6.39, df = 6, *P* = 0.38), with 55% to 83% of egg masses in average giving rise to a larva (**Fig. S1B-C**).

By contrast, we observed a striking difference in the distribution of egg masses between the two strains. For each experimental trial (“choice”, “corn” and “rice”), sf-C laid between 33% to 52% and sf-R laid almost 85% of their egg masses on the cage net rather than on a plant (**Fig. 1**). Neither strain showed a preference for the expected host-plant in female’s oviposition choice (*i.e.* corn for sf-C and rice for sf-R). Behavior difference between strains was indicated by the highly significant interaction between strain and laying site in all trials (LRT for maize trial: χ^2^ = −68.35, df = 1, *P* < 0.001; LRT for rice trial: χ^2^ = −90.10, df = 1, *P* < 0.001.; LRT for choice trial:χ^2^ = −39.53, df = 2, *P* < 0.001). For sf-C, our model shows no difference in the proportion of egg masses between the net and corn plants in corn trial (LRT, χ^2^ = −1.30, df = 1, *P* = 0.25) but did show a significantly (LRT, χ^2^ = − 20.03, df = 1, *P* < 0.001) higher number of egg masses on rice plants than on the net in rice trials (**Fig. 1A-B**). For sf-R, in the no choice trial, the females laid more eggs on the net than on plants (LRT for maize trial: χ^2^ = −83.99, df = 1, *P* < 0.001; LRT for rice trial: χ^2^ = − 72.95, df = 1, *P* < 0.001; **Fig. 1C-D**). In the choice trial, both strains exhibited the same preference pattern. Indeed, the proportion of egg masses for both strains was higher on the net than on corn (sf-C strain: χ^2^= −8.2766, df = 1, *P* < 0.01; LRT for sf-R strain: χ^2^ = −60.65, df = 1, *P* < 0.001) or on rice (sf-C strain: χ^2^= −44.949, df = 1, *P* < 0.001; LRT for sf-R strain:χ^2^ = −98.30, df = 1, *P* < 0.001) and lower proportions on rice than on corn (sf-C strain:χ^2^= −15.23, df = 1, *P* < 0.001; sf-R strain:χ^2^= −7.28, df = 1, *P* < 0.01; **Fig. 1E-F**).

**Figure 1:**
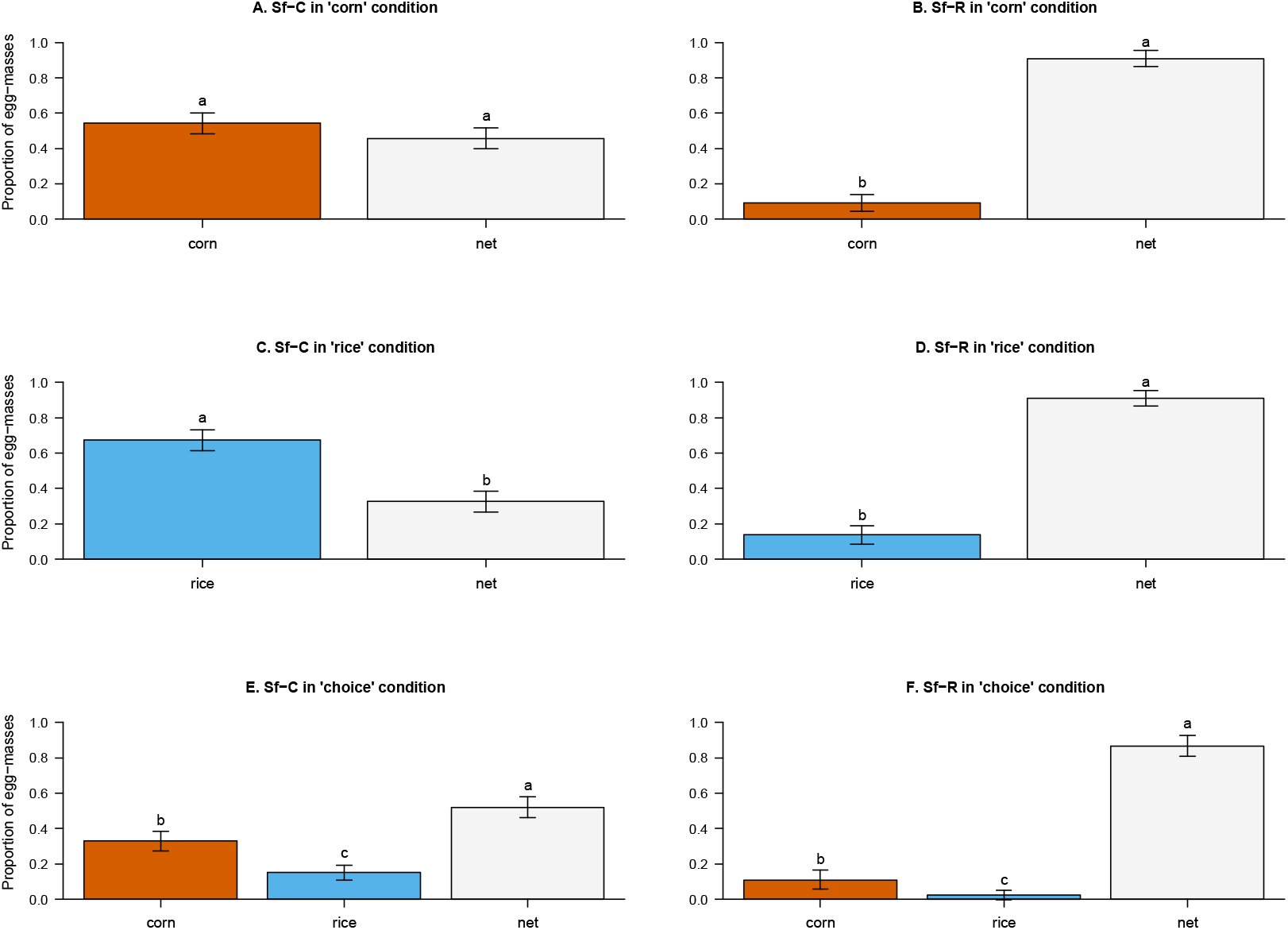
oviposition choice of sf-C and sf-R. Proportion of egg masses laid in the three experimental trials (corn-only, rice-only and choice) by sf-C **(A-C-E)** and sf-R **(B-D-F)** according to the site of oviposition. There are three oviposition sites available: the net (light gray), the corn plant (red) and the rice plant (blue). Here, the relative proportions on each laying site represented the mean of proportions obtained about the four replicates.

While these results did not detect a plant host preference for egg laying, behavioral differences between strains were observed, with sf-C laying more egg masses than sf-R, and sf-R placing more egg masses on the cage surface than on plants. This lack of preference for their preferred host plant is surprising because *S. frugiperda* is a species subdivided into two strains according to the host plant on which the individuals were found preferentially (*i.e*. sf-R on *Oryza sativa*, *Cynodon spp*. (including the Bermuda *grass Cynodon dactylum*) and the legume *Medicago sativa* (alfalfa) whereas sf-C consumes mainly *Zea mays*, *Sorghum* spp. and *Gossypium hirsutum*; Pashley 1986). The question of qualifying them as two distinct species has already been raised (Dumas et al. 2015). However, although two variants are defined, *S. frugiperda* is mainly qualified as a polyphagous species found on more than 353 different host plants belonging to 76 different families (Montezano et al. 2018). Despite these host plant preferences observed in natural populations, both strains can be sampled on the same plants (Juárez et al. 2012). About 19% of sf-R individuals are present on maize and 5% of sf-C individuals are present on various herbaceous plants (Prowell et al. 2004). This lack of striking female preferences could be accentuated by working on laboratory strains, forced for several generations to lay on filter paper.

### Larval fitness in RT experiment

To test whether different plant diets have an effect on the fitness of *S. frugiperda* larvae, we performed a series of reciprocal transplant (RT) experiments in which larvae freshly hatched of both strains were deposited in cages containing either their current or their alternative host plant. Larvae were allowed to develop on their plants, with the food source being regularly supplied as to avoid deprivation. A control population was reared in parallel on the “Poitout” artificial diet normally used to culture the insects in the laboratory (Poitout and Bues 1974). During the experiment, we recorded several phenotypic traits: the weight (wt), the developmental stage to measure the time intervals (dt) and the survival (sv).

After hatching, *S. frugiperda* larvae of the first stage (L1) have to undergo five molts to reach their 6^th^ and final stage (L6) prior to metamorphosis. The time intervals between stage (dt) was explained only by the host plant (LRT, χ^2^ = −37.41, df = 1, *P* < 0.001) and there was no strain effect (LRT, χ^2^ = −0.93, df = 1, *P* = 0.335; **Fig. S2E-F**). In sf-C, the larvae took about 11 to 12 days to complete their larval cycle feeding on artificial diet. We obtained the same duration (11 days) with larvae feeding on corn. Remarkably, development of sf-C larvae feeding on rice took 6 to 7 days longer compared to the other diets (**Fig. S2E**). The sf-R larvae took 11 to 13 days after hatching to complete their larval development on corn compared to 17 days for artificial diet and rice (**Fig. S2F**). Finally, both strains exhibited a similar pattern for dt from 1^st^ larval instar to adult emergence, with both strains having a longer dt feeding on rice than on corn (LRT, F = 28.88, df = 1, *P* < 0.0001; **Fig. S2E-F**). Development on corn was similar for both strains (17 days), but sf-R grew faster on rice than sf-C (22 against 24 days, LRT: F = 182.38, df = 1, *P* < 0.0001).

Weight (wt) at the pupal stage was explained by host plant (LRT: χ^2^ = −555.25, df = 1, *P* < 0.001), moth strain and sex, with a significant interaction between the last two variables (LRT: χ^2^ = − 6.61, df = 1, *P* = 0.012). Indeed, we observed, except for sf-C on corn, that males were heavier than females (**Fig. 2**). Both strains had heavier pupae from feeding on corn than feeding on rice (for sf-C: LRT, χ^2^ = 67.107, df = 2, *P* < 0.001; for sf-R: LRT, χ^2^ =27.18, df = 2, *P* < 0.0001, **Fig. 2**). Pupal weights were higher on corn condition (around 260 mg) than on rice (around 185 mg; **Fig. 2A-C**). Overall, sf-R larvae and pupae were much lighter than sf-C larvae. In all feeding regimes, the maximum larval weight was between 260 mg and 410 mg, while the pupal weight was between 115 mg and 180 mg. Larvae did best feeding on corn, with higher weight gain than on the artificial diet or on rice (**Fig. 2B-D**).

**Figure 2:**
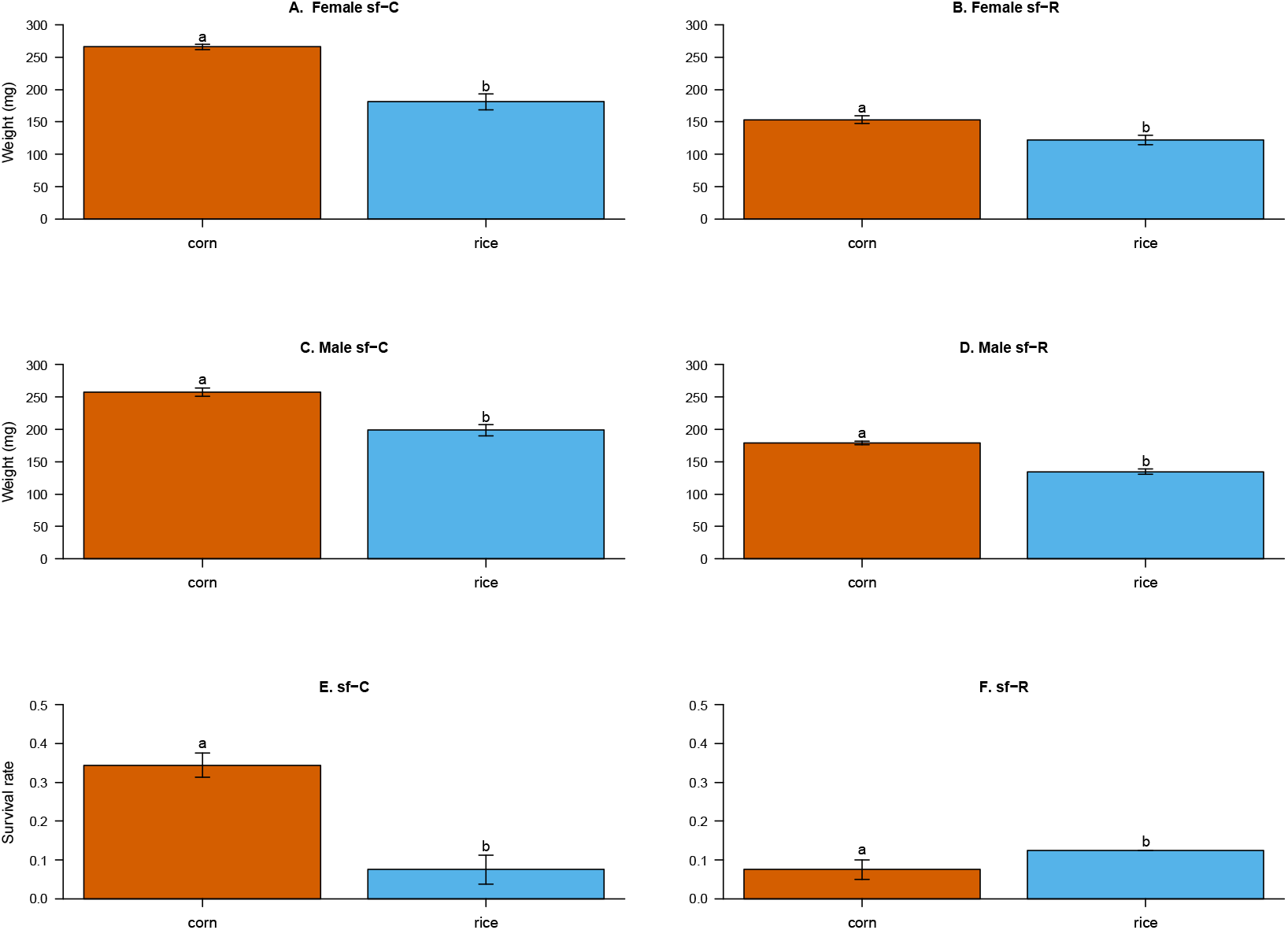
Fitness traits of sf-C and sf-R according to the diet. **(A-D)** Pupal weight (wt) is measured in duplicate for sf-C **(A-C)** and sf-R **(B-D)** according to plant diet: corn (red) and rice (blue). We measured separately females **(A-B)** and male **(C-D)** pupae. The survival (sv) rate **(E, F)** is measured from the 1st larval instar to adult emergence for sf-C **(E)** and sf-R **(F)** according to plant diet. Bars represent the mean of survival rate of the two experimental replicates with the standard error. Different letters above bars indicate significant differences of survival between plant diets for each strain (P < 0.05).

The survival (sv) of both strains was linked to the host plant on which the larvae developed. There was a significant interaction to sv between strain and host plant (LRT, χ^2^ = −24.22, df = 1, *P* < 0.0001; **Fig. 2C-D**). The survival of sf-C was significantly greater on corn (about 34%) than on rice (about 7.5%; **Fig. 2C**). However, although sf-R tended to have higher survival on rice (LRT, χ^2^ = 2.53, *P* = 0.11), sv was not significantly different between the two host plants (7.5% on “corn” vs 12.5% on “rice”; **Fig. 2D**). We noted that the survival rates on plant experimental set-ups were relatively low, with most mortality occurring during larval stages. These absolute numbers cannot be related to controlled conditions where artificial rearing is designed to provide as much survival of the population as possible (**Figure S3**). Similarly, it cannot be compared to survival rates in the wild, for which we have no estimate. Host-plant, but also variable environmental parameters and interactions with competitors, predators, parasites and pathogens can affect the survival and are an essential component of the host-plant as an ecological niche. Here, we can only conclude on the relative survival rates between similar experimental conditions, which we think reveals intrinsic adaptation to the host-plant.

In brief, this analysis indicates that under our laboratory conditions, there is a clear effect of the host plant on the fitness of *S. frugiperda*. Individuals of both strains grew faster and gained more weight feeding on corn than on rice. We observed one major difference between strains, with sf-C surviving better on corn than sf-R, suggesting a specialization of sf-C to corn. However, we didn’t find the reciprocal trend for sf-R, which survived equally on both plants. Once again, as noted in the plant preference, the absence of plant cues during laboratory breeding over several generations could have allowed a relaxed selection of host plant characteristics. Moreover, the artificial diet is based on corn flour and therefore Sf-R has not been confronted with rice compounds for many years, which might explain why differences between these two plants are not detected.

### Gene expression in RT experiment

When confronted with different host plants, polyphagous insects will respond by expressing different sets of genes, some of them can be associated to a better adaptation to the host plant. Such adaptation genes in insects are known to be involved in chemosensory, digestion, detoxification and immunity processes among others (Simon et al. 2015; Celorio-Mancera et al. 2016). In order to understand if the two *S. frugiperda* strains express different adaptation genes to host plant diet, we performed RNA-Seq experiments from the larvae of the RT experiments. RNA was extracted from 4^th^ instar larvae from the same RT experimental setup as the one on which LHT were measured. We could perform for each strain two replicates on the corn diet, one replicate for the rice diet and one replicate for the artificial diet. We recovered between 30 to 71 million reads per sample (**Table S1**), which we aligned on the OGS2.2 reference transcriptome for sf-C (Gouin et al. 2017) containing 21,778 sequences. The percentages of reads mapped were similar between the two moth strains, with 72.1% to 73.3% of alignments for sf-C under any diet (**Table S1**). For sf-R samples on corn the alignment percentages were similar (71% and 71.2%), and slightly less for the other samples (68.6% on artificial diet and 68.9% on rice; **Table S1**). Hereafter, we refer to this RNA-Seq dataset as MORT2.

### Constitutive transcriptional differences between sf-C and sf-R

PCA analysis of the MORT2 RNA-Seq data shows that the samples are grouped by strain (29% of explained variance on PC2; **Fig. 3A**), suggesting there may be fundamental differences between sf-C and sf-R that could explain their plant preferences. However, this observation was contrasted by PC1, which explained 53% of the variance and revealed a pattern of separation by preferred diets. Indeed, an important part of the variance was explained by the sample sf-R on rice, clustering with sf-C on corn (**Fig. 3A**). We used DESeq2 (Love, Anders, and Huber 2014) to identify constitutive differences between the two strains regardless of the diet trial. We identified 1,697 (7.8%; *p.adj* < 0.05) genes overexpressed in sf-R compared to sf-C and 2,016 (9.3%; *p.adj* < 0.05) genes overexpressed in sf-C compared to sf-R (**Fig. 3B**). We verified by q-PCR on independent samples raised on artificial diet that this strain-specific difference of expression is stable. We selected and annotated (**Fig. S4**) 50 genes overexpressed in sf-R compared to sf-C in our RNA-Seq experiments (**Fig. S5-S6**), all except one (peroxidase), were systematically overexpressed in sf-R when measured by qPCR (**Table S2-S3 & Fig. S7**).

**Figure 3:**
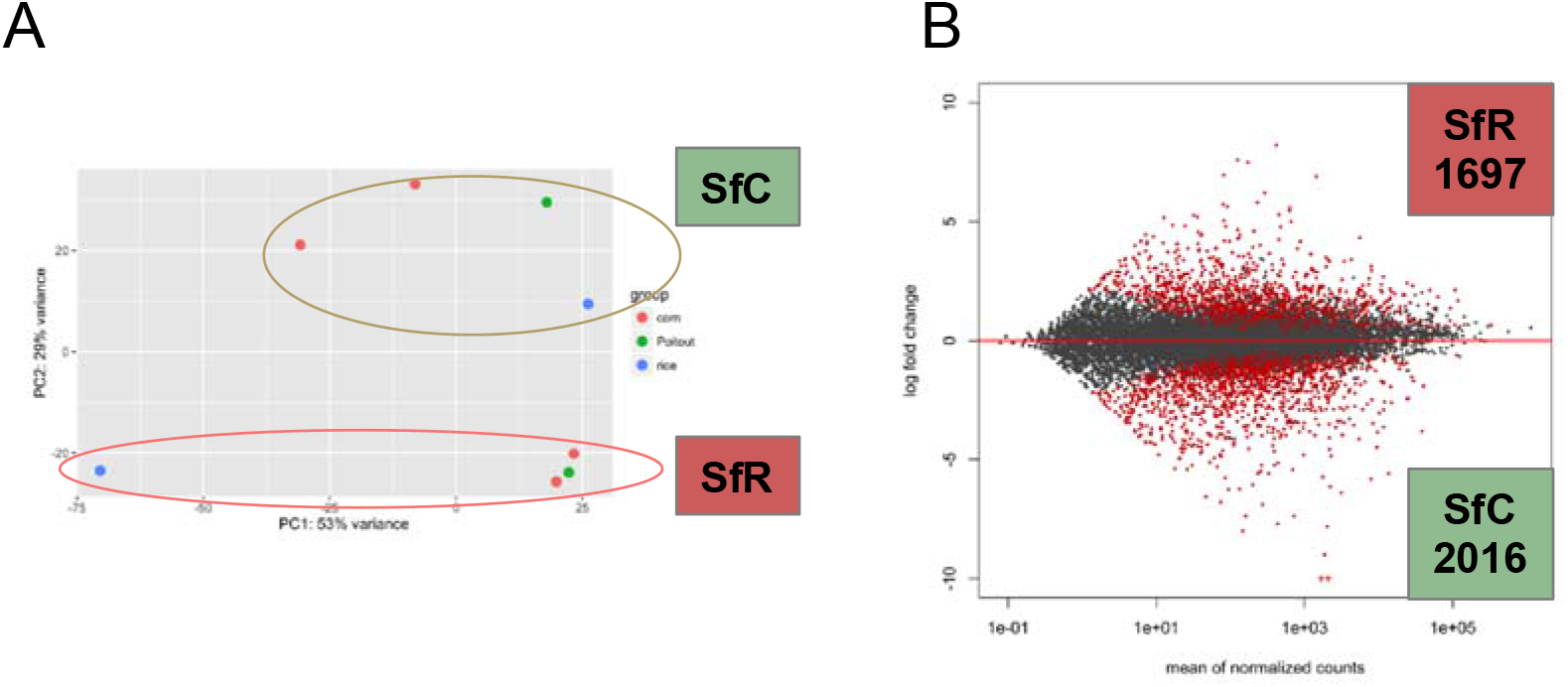
Transcriptional response of sf-R versus sf-C regardless of the diet. **(A)** Principal component analysis on normalized RNA-seq reads for all RT samples of sf-R and sf-C when the larvae feed on corn (red), on rice (blue) or on artificial diet (green). **(B)** Multidimensional scaling plot (MA-plot) reporting the log2 fold changes between the strains (sf-R vs sf-C) over the mean of normalized counts. Each dot represents a gene either with a non-significant differential expression between trials (gray dots) or with a significant differential of expression (red dots).

The GO enrichment analysis did not detect any significant enrichment of either Biological Process or Molecular Function terms in both gene lists. sf-R expresses some enzymes involved in digestion, metabolism and detoxification as well as, intriguingly, ribonucleoproteins involved in mRNA splicing (**Fig. S5**) but no coherent pattern emerges. While no GO enrichment has been observed for sf-C, manual re-annotation of the 50 most expressed genes showed that at least 13/50 genes correspond to transposable elements (TE) (**Fig. S6**). Other genes encode putative endonucleases that could also be of TE origin, such as the Harbinger transposase-derived nuclease, HARBI. In addition, we could not find evidence for gene annotation by homology or protein domain analysis for 16/50 genes. Other genes encode proteins that could be linked to plant adaptation. For example, sf-C shows a strong expression of fatty acid synthase, suggesting that sf-C is constitutively more efficient at energy production and storage. We also found two peptidases, and the cytochrome P450: CYP9A31 indicating inherent digestive and detoxification potential for sf-C. While we have detected no transcriptional regulators in our plant adaptation datasets, we could at this time detect one important transcription factor (TF), expressed only in sf-C: *apterous-1*. This homeodomain (HD)-containing TF is known in *Drosophila* to be involved in wing development (http://flybase.org/reports/FBgn0267978.html). Annotation of HD genes in *Spodoptera* (Gouin et al. 2017) showed that *apterous* has two paralogs, suggesting a yet-to-be-determined potential shift in function for this TF. Finally, we detected overexpression of a small genomic sequence corresponding to a fragment of the mitochondrial gene cytochrome oxidase c subunit III (COIII). Genomes often contain insertions of mitochondrial sequences (Hazkani-Covo, Zeller, and Martin 2010). Such insertions are termed *numts*. Around 95 *numts* can be identified in the *S. frugiperda* genomes. They sometimes confound gene prediction because they contain the open reading frame (ORF) sequence of the original mitochondrial gene. However, *numts* are usually not transcribed, lacking the promoter region sequence and the measured differential expression comes from messenger RNAs of mitochondrial origin, whose reads also align on the numt region (**Fig. S16**). Thus, in practice, *numts* can be used to measure the expression of portions of the mitochondrial genome.

### Exploration of strain transcriptional differences in natural populations

We wanted to know if the transcriptional differences between *S. frugiperda* strains measured in the laboratory conditions can also be observed in the wild. We performed a field collection of FAW larvae in a sweet corn field (Citra, FL), in a volunteer corn field (Tifton, GA) and in a pasture grass field (Jacksonville, FL). We performed both DNA and RNA extractions from individual L4 larvae. DNA was used to genotype the individuals (see **Methods**). Based on the detection of mitochondrial Cytochrome Oxidase I (COI) polymorphism (Nagoshi et al. 2006), the Citra corn field contained 32/33 sf-C associated genotypes, the Tifton corn field contained 14/18 sf-C strains and the Jacksonville field contained 6/6 sf-R strains (**Fig. S8**). We selected some sf-R and sf-C individuals from each field to genotype according to one SNP on the *Tpi* gene located on the Z chromosome (Nagoshi 2010) and presence of the FR1 repeat (Lu et al. 1994; Nagoshi and Meagher 2003a). Interestingly, most sf-R haplotypes recovered from corn fields seem to be hybrids from a sf-R mother. We didn’t detect any potential hybrids in the pasture grass field (**Fig. S9-S10**).

From the 20 most differentially expressed genes between sf-C and sf-R on corn, we selected 15 genes to perform qPCR measurements of their expression in individual L4 larvae from the laboratory strains raised on the artificial diet as well as in individual L4 larvae from the Tifton field where we recovered both sf-C and sf-R mitochondrial haplotypes. The qPCR analysis showed that the genes we selected from RNA-Seq studies are concordantly differentially expressed between laboratory strains. However, for the genes we selected, we detected no difference in expression between natural populations of sf-C and sf-R (**Fig. S11**). This result seems to indicate that studies of plant adaptation in laboratory conditions might not be directly applicable to natural conditions. Indeed, in laboratory conditions, we can control the genetic background of insects, the environmental conditions as well as the plant types and supply, while natural populations experience many more variables. Their genetic background might be different from one another, they may be infected or parasitized, they may be individually stressed by climate conditions, predators, competitors or parasitoids. In these conditions, to identify transcriptional differences between strains, one might want to turn to RNA-Seq experiments, which allow interrogating all genes at once.

### Transcriptomic studies of natural *S. frugiperda* populations

We thus decided to produce a dataset (named FL15 hereafter) of RNA-Seq experiments with 3 sf-C individuals from Tifton, 3 sf-R individuals from Tifton and 3 sf-R individuals from Jacksonville (**Fig. 4A**). We recovered from 23 to 74 million reads per sample (**Table S1**) with alignment percentages ranging from 45.32% to 58.40%, slightly less than in laboratory experiments. On a PCA analysis of FL15 dataset only, replicates of the same “trial + strain” individuals group well together with the FL15_B1J individual being slightly outlier (**Fig. S12A-B**). When integrating all FL15, and RT experiments, it becomes impossible to group together all Sf-C genotypes independently of trials (**Fig. S12C**). Moreover, when we looked at the expression of the 50 most differentially expressed genes in sf-R versus sf-C in RT2 experiments and observed the expression of these genes in two independent RT experiments RNA-Seq from our laboratory (RT1), a previously published study on the midgut Roy-RT (Roy et al. 2016) and the FL15 natural populations, we observed that most transcriptional response detected in RT2 was not recapitulated in the other experiments (**Figs. S13-S14**).

**Figure 4:**
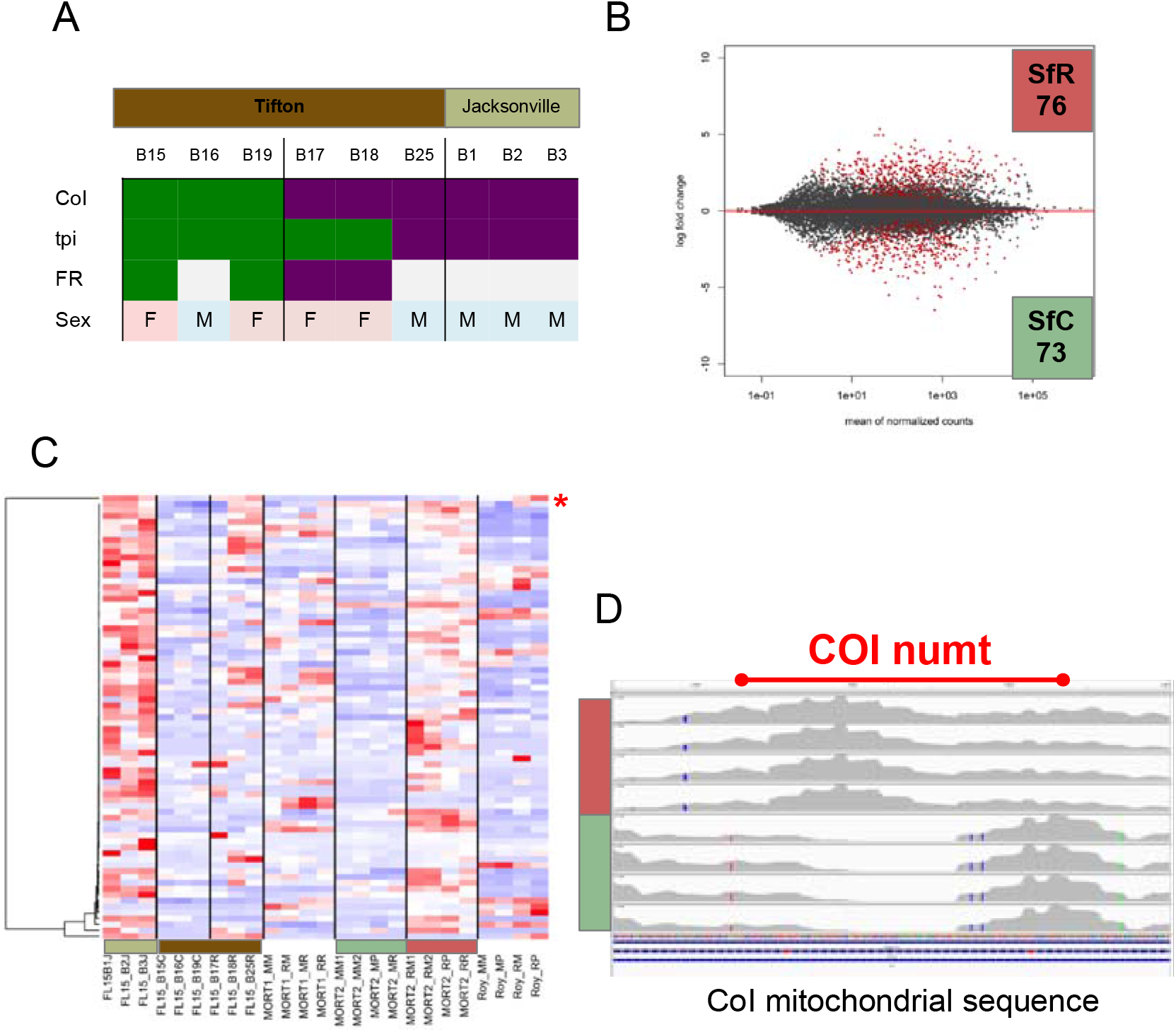
RNA-Seq of individual larvae from the fields. **(A)** Genotypes of the individual L4 larvae from natural populations used for RNAseq studies. CoI, tpi and FR1 repeat genotyping has been done by PCR-RFLP (Supplementary Figures 8-10). Color code is dark green for presumptive C-strain genotype according to the literature while purple is for presumptive R-strain genotypes. Sex has been determined post-facto by examining the alignments of reads on the Z-associated tpi locus. If all SNP positions within the scaffold are homozygous, we assumed the individual was female. Heterozygosity indicates a male. **(B)** Multidimensional scaling plot (MA-plot) reporting the log2 fold changes between the strains (sf-R vs sf-C) over the mean of normalized counts when combining FL15 and MORT2 experiments. Each dot represents a gene either with a non-significant differential expression between conditions (gray dots) or with a significant differential of expression (red dots). 76 genes are overexpressed in sf-R and 73 in sf-C. **(C)** Heatmap of expression variations (expressed as z-scores) of the sf-R specific expressed genes across all RNAseq experiments. For each gene, red indicates a higher expression and blue a lesser expression across the experimental dataset. Genes have been hierarchically clustered as indicated by the dendrogram on the left by similarity of expression variation. The red asterisk identifies the COI-numt expression. **(D)** View of the mitochondrial genome corresponding to the COI-numt sequence and alignment coverage of reads corresponding to sf-R (red) or sf-C (green) samples of the MORT2 experiment. We can observe a trough of expression in this region associated with sf-C strain.

### Strain specific expression in laboratory and in field collections

We took advantage of a large dataset to ask again a simple question: what are the genes whose expression is constitutive of one strain compared to the other? We performed a differential expression analysis across our laboratory RT experiment and our FL15 collection to identify these genes. We found 76 genes consistently overexpressed in sf-R compared to sf-C and 73 genes overexpressed in sf-C compared to sf-R (**Fig. 4B**). To verify the validity of these genes we again surveyed their expression across all the RT-RNA-Seq data at our disposal. We could see that for the majority of these genes their strain specific overexpression is confirmed in the different laboratory populations as well as in natural populations (**Fig. 4C** and **Fig. S15**). Many genes in this list have functions of potential interest to study the molecular basis of ecological speciation (**Tables S4 & S5**). As noted with laboratory sample RNAseq experiments, the sf-C associated overexpression points to many genes whose manual annotation reveal transposable elements of the PiggyBac and Ty1/Copia families (**Table S4**) suggesting a recent reactivation of transposition events in this strain. We also note many genes that could be linked to plant adaptation such as fatty-acyl CoA reductase, OBP36, glucose dehydrogenase, fatty acid synthase, cytochrome P450 and glyoxalase, as well as immunity genes such as a GNBP, a lectin and a cecropin. Finally, this list comprises some potential regulators of expression such as the homeobox transcription factor apterous-1, the DNA helicase Pif1, Orc4, the Mcm complex, a HMG box factor, NUP62 and Leo1. The genes associated to sf-R overexpression (**Table S5**) have a wider array of function but, interestingly, some members in this list also have the same molecular functions as the sf-C expressed such as Fatty acyl-CoA reductase and Glucose dehydrogenase. We also noted a strong pathway of hormonal regulation with the overexpression of the ecdysteroid kinase and the broad complex, as well as the takeout gene which is a juvenile hormone binding protein involved in foraging behavior in Drosophila and NGFI-A-binding protein co-factor, involved in neuron regulation.

To verify the validity of this gene list, we applied a hierarchical clustering analysis of their expression across all the RNAseq data at our disposal. We noticed peculiar outliers with strong expression associated to sf-R corresponding to the previously mentioned *numts* (**Fig 4C**, **S16**). As mentioned, these *numts* reveal parts of the mitochondrial genome that are differentially expressed according to the strain. Two of these *numts* correspond to fragments of the mitochondrial genes COI and COIII (**Fig. S16**). To rule out any effect of genome misassembly, we amplified both *numts* and mitochondrial sequence for COI and COIII and sequence them. We could confirm the presence of these *numts* within the genome of sf-C and sf-R strains with a sequence slightly different than the one from mitochondria. To rule out any sequence specific alignment bias, we retrieve from NCBI the reference genome sequence from *S. frugiperda* mitochondrion (accession KM362176.1) and realigned our RNA-Seq data on it. It was obvious that, in the regions corresponding to *numts*, there was a clear underexpression in the sf-C strain (**Fig. 4D**). The implication of this result on the metabolism of the larvae remains to be established, but nevertheless, it may explain why the mitochondrial haplotypes in the COI gene is the principal marker for strain discrimination. It may very well be that a difference in energy production between these two strains was linked at some point of their evolutionary history to a shift in host plant preference.

## Conclusion

In this study, we wanted to determine if the differentiation of *S. frugiperda* in two strains - sf-C and sf-R - is a result of their adaptation to different host plant diet. First, we measured a combination of Life History Traits in the context of an oviposition preference experiment (OV) and of a reciprocal transplant (RT) experiment in controlled environments to characterize the specialization to host plants. Then we performed RNA-Seq measurements of gene expression variations of L4 larvae during controlled RT experiments in the laboratory and in natural populations. The integration of these datasets allowed us to reveal constitutive differences between sf-C and sf-R.

From this set of experiments, we concluded that the LHT of our laboratory colonies are consistent with a specialization of sf-C to corn, but does not provide evidence that rice is the preferred plant for sf-R, which showed only a slight trend to survive better on this plant than on corn. Interestingly, however, RNA-Seq experiments show that both strains express a similar set of genes, involved in growth and nutriment storage, when confronted to their main host-plant (corn for sf-C and rice for sf-R). This similarity in the transcriptional responses suggests that rice is indeed recognized as a suitable host for sf-R but maybe not its most preferred one.

We found several candidate genes that are differentially expressed between the strains regardless of the diet. However, when we looked at natural populations, almost none of these genes were differentially expressed between strains. But by combining the analysis of RNA-Seq data from laboratory populations as well as from natural populations, we detected a narrower set of genes constitutively differentially expressed between strains. Among those, one candidate stood out and turned out to be the mitochondrial gene COI. This gene is used as a genetic marker for strain identification in all fall armyworm related publications, including the survey of invasive populations in Africa (Nagoshi et al. 2018). The fact that it is also constitutively differentially expressed may indicate that the COI gene, and potentially other mitochondrial genes, may be the original target of selection between the strains (Meiklejohn, Montooth, and Rand 2007). Changes in mitochondrial functions are associated to changes in energy demand or supply (Jose et al. 2013). In addition, variations in mitochondrial sequences can be the cause of mitonuclear incompatibilities between species (Hill 2015, Ghiselli and Milani 2019). The evolution of mitonuclear interactions can maintain the segregation of various mitochondrial haplotypes in the context of ecological speciation (Morales et al. 2016). These features are consistent with a model of ecological speciation for *S. frugiperda*, in which divergence in mitochondrial functions have been selected on plants with different nutritive values. For example, the sf-C haplotype, which has a lesser expression of mitochondrial genes might have a reduced energy production efficiency compared to sf-R. This reduced efficiency may be compensated by the higher nutritive value of the corn plant. Consistent with this explanation, we found sf-R haplotypes in corn fields but almost no sf-C haplotypes on pasture grass fields. Alternative explanations might involve adaptation to the redox state imposed by the host-plant xenobiotic compounds. Several insect proteins such as UGTs and P450s catalyze oxidation-reduction reactions to resist against these natural pesticides. Consistent with this second hypothesis, we also detected plastic and evolved differential expression of several P450 proteins. Finally, it is possible that variations in mitochondrial function reflect variations in energy demand associated with the different field environments. Indeed, corn plants, especially the hideouts within the whorl or the ear, may also provide more protection against competitors, predators and parasitoids than grass lands, which are more open spaces. Thus sf-R strain, that has a higher level of expression in mitochondrial genes might require more energy to move around. Consistent with this explanation, sf-R larvae are consistently smaller than sf-C larvae (**Fig. 2A-D**). Energy consumptions at the adult stage, especially regarding migratory capacities should also be considered.

Compared to other studies using a similar RT experimental design to identify adaptation genes or evolved genes in *S. frugiperda*, our study highlighted one important point that could explain the inconsistencies observed over the years in the determination of the plant adaptation process in *S. frugiperda*. Traditionally, two different RT strategies were used, either by using colonies from natural populations or long maintained laboratory colonies and each approach has its pros and cons. Working with laboratory colonies allows one to control for genetic background variations as well as environmental conditions. But in turn, they might be subject to genetic drift or adaptation to the artificial diet used to maintain them. Here, we show that by combining the two approaches, we revealed a smaller set of genetic events that could explain the differentiation of the two strains. In particular, we identified COI as both a genetic marker and a functionally different locus between the two strains. The consequences of functional variations in the mitochondrial genome on the shift of host-plant range in *S. frugiperda* remains to be elucidated.

## Material and Methods

### Biological material: Moths and Plants

We used individuals from the two strains of *S. frugiperda*: corn (sf-C) and rice strain (sf-R). Those strains were seeded with around 50 pupae sampled in Guadeloupe in 2001 for sf-C and in Florida (Hardee County) in 2012 for sf-R. From the time of their collection they have been reared under laboratory conditions on artificial diet (from Poitout et al. 1972, principal components: 77% H_2_O, 2% Agar-agar, 13% maize flour, 6% other nutrients, 1% vitamins; 1% antibiotics), at 24°C with a 16h:8h Light:Dark photoperiod (L:D) and 70 % Relative Humidity (R:H).

Corn (Corn line B73) and rice (Arelate variety from CFR, Centre Français du Riz) were produced from organic seed at the DIASCOPE experimental research station (INRAE, Mauguio, France, 43°36′37″N, 3°58′35″E) in plastic pots (7 × 8cm for both plants in RT and 6L plastic pots for maize in OV) filled with conventional substrate. Corn and rice cultivation were carried out in a warm chamber at 25°C 2, 60% RH and 16:8 h (L:D) under organic conditions. Corn and rice plants were used 15 days or a month after seeding, respectively, to have an equivalent of two biomass plants.

### Experimentation

#### Experimental trials

*S. frugiperda* is not yet present in France and considered as a quarantine pest. Consequently, experiments on this organism are regulated. Our experiments described hereafter were conducted in confined environment on an insect quarantine platform (PIQ, University of Montpellier, DGIMI laboratory).

#### Oviposition experiment

The oviposition (OV) experiment consisted in release of 12 to 20 virgin females and males of the same strain per cage, and for three nights (72 hours) in three different set-ups: *choice*, *corn-only* and *rice-only*. All individuals released had emerged the night before the oviposition choice experiment. For the choice modality, each cage contained five maize plants and 15 rice plants (the number of maize and rice were adjusted to provide an equivalent biomass) arranged in two patches in two opposite corners of the cage. For the rice- and corn-only modalities, we used either 10 maize or 30 rice plants. Plants were arranged in two equal patches (2 × 5 maize or 2 × 15 rice) located in two opposite corners of each cage. The experiment was conducted in insect rearing cages covered by an insect-proof knitted-mesh of nylon (175 × 175 cm) and 4 replicates of each set-up were done under the same climatic conditions, within the quarantine platform (22°C, 50% humidity, natural dark-light conditions - in November around 14h dark:10h light-with fluorescent light bulbs).

In each cage, at the end of the third night, all egg masses were counted and immediately individualized. We measured three variables for each cage:

1. The number of egg masses laid by females in a given cage (on plants and on the net) to measure the fecundity. As the adult number was not similar in cages, it was important to balance the number of egg masses per the number of females in the cage. Indeed, the number of adults had a significant effect on the egg masses number (*P* < 0.01), so we decided to create a variable, *Mean Fecundity*, which took into account the egg masses number divided by the number of females in the replicate. The following variables were the strain (sf-C and sf-R) and the trials (choice, rice-only, corn-only).
2. The proportion of egg masses laid by females on one particular site (one given plant species or the net). This percentage was calculated in three set-ups to estimate the preference of each moth host strain according to present substrates in the cage. We performed the analysis on each set-up independently with two following factors, the strain and the oviposition site.
3. The hatching proportion is the number of egg masses hatching on one particular site (one given plant species or the net) whatever the set-up. This percentage provides an estimate of the fertility of both strains according to the choice of oviposition site by the females. The following factors are the strain and the oviposition site (nested in set-up).

#### Reciprocal transplant experiment

The reciprocal transplant (RT) experiment consisted in controlled infestations of corn and rice plants with first instar larvae in 8 insect rearing cages (32.5 × 32.5 cm) covered by an insect-proof veil to prevent contaminations and escapes in the incubator (24°C, 16h:8h L:D cycle and 70% R. H.). The RT experiment was conducted in the same incubator for four modalities: 1) corn plants infested by sf-C (native condition); 2) rice plants infested by sf-C (alternative condition); 3) corn plants infested by sf-R (alternative condition); 4) rice plants infested by sf-R (native condition). We realized two replicates by modality. Each cage contained four corn or rice pots, which were changed before the 4^th^ larval instar and each day after this instar until the pupation.

From a batch of eggs reared on artificial diet, we subdivided the progeny on the three different diets (corn plant, rice plant and artificial diet). A total of 80 larvae (which hatched the morning of the experiment) were deposited in each cage.

Two generations have been conducted on plants; during the first generation we measured life history traits (LHT) for each strain in native and alternative conditions and during the second generation, the larvae had been sampled at 4^th^ larval instar for RNA-Seq experiments.

As of the 2^nd^ larval instar, we measured several LHT every other day until pupation, during which we determined the sex of each individual. In addition, at each counting, we determined the larval stage by the width of the head capsule. To limit the possible contamination between strains, we isolated two floors of the incubator with an insect proof net (150 μm) and to avoid a floor and edge effect, rotations between floor were conducted and cages were randomly deposed after counting. We measured three variables:

- Survival (sv) is the number of emerging adults counted over the initial number of larvae;
- Developmental time (dt) is the number of days between the beginning of the experimental start until adult emergence (mean on all emerging adult in same cage);
- The weight (wt) of individual larvae and of individual pupae of each sex in mg. The day of plant infestation, we weighed the pool of 80 larvae. Then, from the 2nd larval stage, the weight was quantified every other day and each larva was individually weighed.

For all variables from RT, we analyzed by following factors: the strain (sf-C or sf-R) and the host plant (corn or rice). Replicate effect was negligible. In parallel, and as a reference point, we performed the same experimental design and measurements on standard rearing conditions on artificial diet (Poitout and Bues 1974). Two replicates of each strain on artificial diet have been set-up from the same batch of L1 larvae from our laboratory strains. Compared to plant conditions, rearing has been performed in a square plastic box with mesh filter for aeration and food supplied *ad libitum*. Since the rearing conditions differ significantly from plant assays, we considered those experiments as reference and not as control.

#### Statistical analysis of LHT

All computations were performed using “lme4” package (Bates et al. 2015) of the R software version 3.0.3. We used different generalized linear models depending on the distribution of the residuals. For all the variables, we analyzed by following factors and we also included the interaction between the following factors. If the replicates had a negligible influence on model outcome, they were not included in the models (using “glm” function), or if the replicates had a significant effect, they were added as a random factor (using “glmer” with replicate factor in random effect). Model selection was performed as follows: the significance of the different terms was tested starting from the higher-order terms using likelihood-ratio-tests (LRT). Non-significant terms (*P* > 0.05) were removed and factor levels of qualitative variables that were not significantly different were grouped (LRT; Crawley 2007).

### Genomic

#### Sample preparation and sequencing

We collected 4^th^ instar larvae of the second generation on native and alternative plants, corresponding to offspring of the larvae used to estimate the different components of fitness (survival, weight and developmental time). The larvae number was variable between experimental set-ups (n = 3 to 12 larvae). Larval instar was determined by the width of the head capsule (**Figure S.17**), if the larvae were considered like 4^th^ instar, three larvae of the same experimental set-up were pooled. We weighed the pools and crushed them in liquid nitrogen to obtain a fine powder, which was placed in TRIzol® Reagent (Invitrogen) and stored at −80°C. After collection of samples in all experimental set-ups, total RNA was extracted using a TRIzol® Reagent, according to the manufacturer’s RNA protocol. To remove contaminating DNA from RNA preparations, we used DNase from TURBO DNA-free™ Kit (Ambion). Bioanalyzer using 1 μl of total RNA from each sub-pool of three larvae permitted to estimate RNA quantity. The ratio of absorbance 260/280 and 260/230 was used to assess the purity of RNA in each sample. The sub-pools of three larvae, having a good quality (between 1.35 and 2) and quantity (>200 ng/μl), were pooled again to obtain samples corresponding to the four experimental set-ups. On the one hand, the samples from rice plant containing only three larvae because of the survival problem on rice for both strains. On the other hand, the samples on artificial diet and on maize contained 12 larvae (*i.e*. 4 sub-pools of 3 larvae).

High throughput sequencing was performed for the pool samples using Illumina technologies to obtain single-end 50-bp reads. Library construction and sequencing were performed by MGX-Montpellier GenomiX (Montpellier, France) on a HiSeq 2000 (Illumina). For each pool, tagged cDNA libraries were generated using the TruSeq Stranded mRNA Sample Preparation Kit (Illumina) following manufacturer’s protocol.

#### Reference and annotation

All RNA-Seq experiments were aligned against a common reference. This reference is OGS2.2 (Gouin et al. 2017), generated from the sequencing and annotation of the C-strain genome. Gene models result from direct ORF prediction, guided by expression data published earlier (Legeai et al. 2014) and the mapping of RNA-Seq reads. Gene models for selected gene families also underwent an expert annotation by manual curators.

#### Differential expression analysis

To identify differentially expressed genes, we first mapped reads on gene prediction using Bowtie2 (Langmead and Salzberg 2012). We chose to use the same reference for both the sf-C and the sf-R strain samples. For read mapping we used “very sensitive” parameter setting in Bowtie2, which allowed searching extensively for the best alignment for each read. Counting of aligned reads number to each gene is produced by SAMtools program (Li et al. 2009). Then to detect the genes differentially expressed we used DESeq2 (R package; Love, Anders, and Huber 2014). To measure gene expression variations between conditions, DESeq2 uses a negative binomial generalized mixed model. The estimates of dispersion and the logarithmic fold-changes incorporate data-driven prior distributions. Genes were considered differentially expressed if they satisfy a false discovery rate lesser than 1%.

#### Characterizing gene function and comparison between two strains

After identifying differentially expressed genes between two strains for the same food resource, we used the Fisher’s exact test (cut-off of FDR < 0.01) to identify GO categories possibly involved in corn specialization. The resulting list of GO-terms may contain redundant categories (*i.e*. there was a parent-child relationship in enriched function or process). We used REVIGO (http://revigo.irb.hr/) that summarizes and regrouped terms by a clustering algorithm based on semantic similarities (Supek et al. 2011). We used the default parameter (“medium”).

#### Natural populations collections

*S. frugiperda* wild larvae were collected in Florida and Georgia between September, 18th and September, 25th 2015 in three different field locations. One sweet corn field in Citra (Marion County, Florida), one volunteer corn in Tifton, (Tift County, Georgia) and one pasture grass field in Jacksonville (Duval County, Florida). In corn fields, plants were cut and larvae collected in situ. In the pasture grass field, collections were made using a sweeping net. After confirming their identification as *S. frugiperda* according to LepIntercept (http://idtools.org/id/leps/lepintercept/frugiperda.html), larvae were placed in individual plastic cups with cut leaves (either corn or grass) as a food source and brought back in a cooler to the laboratory after a few hours of collection. Once in the laboratory, larvae were sorted according to stage. Stages were measured according to the chart in **Fig. S17**, where the width of the cephalic capsule should match the width of the line for each stage. This chart has been determined based on rearing conditions of lab strains in Montpellier and confirmed with a similar chart based on the rearing of lab strains in Gainesville, Florida. L4 larvae were sacrificed with a razor blade and immediately placed individually in a screw-cap 2ml tube containing 1ml of RNAlater (Sigma; R0901).

#### DNA/RNA extractions

Larvae from field collections were placed in a 1.5ml Eppendorf tube with RLT buffer from Qiagen. Individual larvae were ground using a TissueLyser II from Qiagen (Cat No./ID: 85300) using one bead (size 5mm) by tube and processed for dual DNA and RNA extraction using an AllPrep DNA/RNA Mini Kit (50) (Qiagen Cat. 80204).

#### Genotyping

We used the COI genotype described in (Meagher and Gallo-Meagher 2003) to discriminate between the sf-C and the sf-R strains. A PCR on genomic DNA was performed using the following primer sequences (JM-77: ATC ACC TCC ACC TGC AGG ATC and JM-76: GAG CTG AAT TAG GGA CTC CAG G) to amplify a DNA fragment of 550bp corresponding to the mitochondrial cytochrome oxidase c subunit I. The MspI enzyme is used to reveal a polymorphism between the 2 strains. The COI fragment of the C-strain is digested by MspI to produce a 500bp and a 50bp fragment (**Fig. S8A**).

For the Tpi genotyping we used the following primers as described (Nagoshi 2010): *Tpi*-56 F (5’-CAAAATGGGTCGCAAATTCG-3’) and *Tpi*-850gR (5’-AATTTTATTACCTGCTGTGG-3’). Digestion of the PCR product was made with the AvaII enzyme (**Fig. S9A**).

FR1 repeat genotyping was based on PCR amplification only, as described (Nagoshi and Meagher 2003a) with the following primers: FR-c (5’-TCGTGTAAAACGTACTTTCTT-3’), and FR-2 (5’-GACATAGAAGAGCACGTTT-3’). Amplification is then analyzed on agarose gel (**Fig. S10**)

#### Quantitative PCR

For reverse transcription quantitative PCR, we used the candidate transcript sequence, as retrieved from BIPAA platform (https://bipaa.genouest.org/sp/spodoptera_frugiperda_pub/) - for example by searching GSSPFG00029721001-RA from **Table S2** - as a template for primer design using Primer3 and asking for a 50 nt amplicon. Primers used are specified in **Table S3**.

Quantitative PCR have been performed on a LightCycler 480 (Roche) with SYBR green. Program used was 95°C for 10min and then 40 cycles of 94°C 10s, 60°C 10s, 72°C 10s. Relative expression was calculated using the ΔΔCt method with the laboratory sf-C strain as a reference point for each gene.

## Data accessibility

*Spodoptera frugiperda* reference genome and reference transcriptome can be publicly accessed via the BIPAA (BioInformatics Platform for Agroecosystem Arthropods) interface (https://bipaa.genouest.org/sp/spodoptera_frugiperda_pub/). fastq files and RNAseq counts from this study are accessible in ArrayExpress (https://www.ebi.ac.uk/arrayexpress/) with the following accession number: E-MTAB-6540. Life history traits data are available online: Marion Orsucci, & Nicolas Nègre. (2020). Data and scripts [Data set]. http://doi.org/10.24072/pci.evolbiol.100102

## Supplementary material

Supporting information, script and codes are available online: Marion Orsucci, & Nicolas Nègre. (2020). Data and scripts [Data set]. http://doi.org/10.24072/pci.evolbiol.100102

## Acknowledgements

This work was partially supported by funding from Institut Universitaire de France for N.N. and by a grant from the French National Research Agency (ANR-12-BSV7-0004-01; http://www.agence-nationale-recherche.fr/) for E.d’A. including a post-doctoral fellowship for Y.M. We thank the quarantine insect platform (PIQ), member of the Vectopole Sud network, for providing the infrastructure needed for pest insect experimentations. We are also grateful to Clotilde Gibard and Gaëtan Clabots for maintaining the insect collections of the DGIMI laboratory in Montpellier, and Amy Rowley for maintaining the original colonies in Gainesville. Version 2 of this preprint has been peer-reviewed and recommended by Peer Community In Evolutionary Biology (https://doi.org/10.24072/pci.evolbiol.100102).

## Authors contributions

NN, EA and MO designed the project. JPB and MV produced the corn and rice plants used in the RT experiments. MO, PA, NN and performed the RT and OV experiments. MO, GD performed the statistical analyses of LHT in the RT experiments. MO, GD, RNS and NN performed the RT-qPCR experiments. MO performed the RNA extractions for the RNA-Seq experiments. RK and SR produced the Illumina libraries, performed the Illumina sequencing and realized the computational analyses and quality control necessary to produce. fastq files of sequences. MO, YM, SN and NN performed the RNA-Seq analyses. MF, GJK, RNN, RLM and NN performed the field collections. RNS and NN performed the genotyping and RNA extractions of field samples. MO and NN wrote the manuscript and produced the figures. YM, SR, GJK, RNN, RLM and EA edited the current manuscript. All authors approved the present manuscript submission.

